# Synovial Gene Signatures Associated with the Development of Rheumatoid Arthritis in At Risk Individuals: a Prospective Study

**DOI:** 10.1101/2021.04.27.440770

**Authors:** Lisa G.M. van Baarsen, Tineke A. de Jong, Maria J.H. de Hair, Johanna F. Semmelink, Ivy Y. Choi, Danielle M. Gerlag, Paul P. Tak

**Author notes:** Corresponding author: Lisa G.M. van Baarsen.

## Abstract

**Background:** Previous work has shown subtle infiltration of synovial T cells in the absence of overt synovial inflammation in individuals at risk of developing rheumatoid arthritis (RA).

**Objective:** To study the molecular changes in synovium preceding arthritis development in at risk individuals.

**Materials and methods:** We included sixty-seven individuals with arthralgia who were IgM rheumatoid factor (RF) and/or anti-citrullinated protein antibody (ACPA) positive and without any evidence of arthritis. All individuals underwent mini-arthroscopic synovial tissue sampling of a knee joint at baseline and were followed prospectively. An explorative genome-wide transcriptional profiling study was performed on synovial tissue using Agilent arrays (discovery cohort). Survival analysis was used to identify transcripts associated with arthritis after follow up. Expression levels of differentially expressed genes were validated using quantitative real-time PCR (qPCR). Immunohistochemistry was used to study gene candidates at the protein level in situ.

**Results:** In the discovery cohort, 6 of the 13 at risk individuals developed RA after a median follow-up time of 20 months (IQR 2 – 44; pre-RA). The 7 individuals who did not develop RA had a median follow-up time of 85 months (IQR 69 – 86). Using a False Discovery Rate of <5% we found increased expression of 3,151 transcripts correlating with a higher risk of arthritis development, whereas increased expression of 2,437 transcripts correlated with a lower risk. Gene set enrichment analysis revealed that synovial biopsies of pre-RA individuals display higher expression of genes involved in several immune response-related pathways compared with biopsies of individuals who did not develop RA. In contrast, lower expression was observed for genes involved in extracellular matrix receptor interaction, Wnt-mediated signal transduction and lipid metabolism. Two-way hierarchical cluster analysis of 27 genes measured by qPCR classified the synovial biopsies of 61 individuals into two groups, where pre-RA individuals (n=16) showed a preference to cluster together. Synovial tissue from pre-RA individuals were more likely to show podoplanin positive cells and lower lipid staining compared with synovial tissue from individuals who did not develop RA.

**Conclusion:** Molecular changes can be detected in synovial tissues before clinical onset of arthritis. Alterations in the immune response genes and lipid metabolism are associated with development of arthritis.

## Introduction

Rheumatoid arthritis (RA) is a chronic inflammatory autoimmune disease which primarily affects the synovial joints and is characterized by chronic synovial inflammation (1). While current treatment options can suppress synovial inflammation and progression of joint destruction in many patients, continued treatment is usually required to prevent relapse (2–6).The ultimate goal of future treatment should be disease remission without the need for continued treatment, in other words cure. Furthermore, insights into the earliest phases of the disease, before the development of clinical signs and symptoms of arthritis, may facilitate the development of preventative approaches (7, 8).

The earliest stages of autoantibody positive RA are characterized by the presence of RA-specific autoantibodies including rheumatoid factor (RF) and/or anti-citrullinated protein antibodies (ACPA) (9, 10). The autoantibodies can be present in the peripheral blood years before clinical manifestation of the disease (11–13). The presence of these autoantibodies indicates systemic autoimmunity and many autoantibody-positive individuals develop RA after follow up (9, 11–14). It has been shown by detailed studies using both immunohistochemistry and MRI that the presence of circulating autoantibodies precedes overt synovial tissue inflammation during the at risk phase of RA (9, 10). However, it has been suggested that there might be subtle T cell infiltration in the synovium during the at risk phase (10).

It is important to identify additional risk factors beyond RF and ACPA in this multifactorial disease, to be able to improve algorithms predictive of which individual will develop clinically manifest RA over time. Previously recognized risk factors include the presence of dominant B cell receptor clones in peripheral blood (15), musculoskeletal symptoms like arthralgia (16), having been a smoker, obesity (17) and decreased vagus nerve tone (18). In this context, we conducted a prospective study and performed explorative genome-wide transcriptional profiling of synovial tissue obtained during the preclinical stage of RA. In the discovery cohort we compared synovial gene signatures of at risk individuals who later developed RA (pre-RA) with those who did not develop RA after a follow-up time of at least two years. Next we used real-time PCR to validate the expression levels and immunohistochemistry was used to study protein expression in situ. Together, we were able to identify molecular changes in the synovial tissue obtained at baseline associated with arthritis development after follow-up. If confirmed in future independent studies, these synovial data may contribute to the identification of novel targets for preventive treatment.

## Materials and methods

### Study subjects and mini-arthroscopy

Individuals with arthralgia and/or a family history of RA who were positive for IgM-RF and/or ACPAs (detected by the anti–cyclic citrullinated peptide [anti–CCP] antibody test) but without any evidence of arthritis upon thorough physical examination, were included in the study (10). These individuals were considered to be at risk of developing RA, a status characterized by the presence of systemic autoimmunity associated with RA (19) without clinical arthritis (defined as phase c+d, according to EULAR recommendations (19)). IgM-RF was measured as previously described (10). The study subjects were recruited either via the outpatient clinic of the Department of Clinical Immunology and Rheumatology at the Academic Medical Center, Amsterdam, via referral from the rheumatology outpatient clinic of Reade, Amsterdam, or via testing family members of RA patients in the outpatient clinic or at public fairs across the Netherlands. The study was performed according to the principles of the Declaration of Helsinki (20), approved by the Institutional Review Board of the Academic Medical Center and all study subjects gave their written informed consent.

All study subjects underwent mini-arthroscopic synovial tissue sampling of a knee joint at baseline (21). To correct for sampling error, 6–8 synovial tissue samples are analyzed together, as described previously (21). Immediately after collection, synovial tissue samples were snap-frozen for RNA extraction or snap-frozen after embedding in Tissue-Tek OCT compound (Miles) for immunohistochemistry analyses. Samples were stored in liquid nitrogen until further processing.

### RNA extraction

Total RNA was extracted from synovial tissue using RNA STAT-60^™^ (Isotex Diagnostics, Friendswood, TX) followed by a cleanup using the RNeasy Mini kit from Qiagen (Venlo, the Netherlands). First, synovial tissue was quickly homogenized on ice in 1 ml STAT-60^™^ solution using an IKA T10 basic homogenizer (S 10 N 5-G probe; 3304000, Cole-Parmer, USA). The homogenized tissue suspension was transferred to a clean RNAse-free tube and incubated at room temperature (RT) for 5 minutes. After adding 200 μl chloroform, the tube was inverted several times to ensure mixing, rested for 3 minutes at RT and centrifuged for 15 minutes at 12,000 g at 4 degrees Celsius. The resulting upper aqueous layer containing RNA was transferred to a new tube and 500 μl isopropanol was added. After 5-10 minutes incubation at RT, tubes were centrifuged allowing RNA precipitation. After washing the pelleted RNA in 1 ml 75% ethanol, RNA was dissolved in 30 μl RNase-free water and stored until further use at −80 degrees Celsius. For RNA cleanup the remaining RNA volume was adjusted to a total volume of 100 μl in RNase free water after which 350 μl of RTL buffer from the Qiagen kit was added and mixed by pipetting. Next, 250 μl 96-100% ethanol was added and sample was mixed by pipetting before transferring the sample to a RNeasy Mini spin column. Further RNA extraction was performed according to the manufacturer’s instructions including an on-column DNase digestion using the RNase-Free DNase Set (Qiagen). RNA purity and quantity was measured using the Nanodrop (Nanodrop Technologies, Wilmington, USA) with ND1000 V3.8.1 software (ND1000, Isogen Life Science, Utrecht, the Netherlands). Cleaned RNA was stored at −80 degrees Celsius until further use.

### Microarray analyses

T7-based linear RNA amplification, Cy-dye labelling and hybridization to Agilent High Definition 4×44k array (Agilent technologies, Amstelveen, the Netherlands) was performed as described previously (22). Samples were analyzed against the Stratagene Universal Human Reference (Agilent). Raw fluorescence intensities were quantified and normalized (Lowess normalization) using Agilent Feature Extraction software v9.5 (Agilent, Santa Clara, CA) according to the manufacturer’s protocols. Log10-ratios (sample/reference) were used for downstream statistical analyses.

Survival analysis within the software package Significance Analysis of Microarrays (SAM) was used to identify transcripts with a significant association with arthritis development (23). Cluster analysis was used to define clusters of coordinately expressed genes and to classify patients according to their transcriptome profile (24). Cluster diagrams were visualized using Treeview (JAM software GmbH, Trier, Germany).

### Quantitative real-time PCR

RNA (500 ng) was reverse transcribed into cDNA using the RevertAid H-minus cDNA synthesis kit (MBI Fermentas, St. Leon-Rot, Germany) according to the manufacturers’ instructions. Quantitative real-time PCR (qPCR) was performed using a Step One Plus detection system (Applied Biosystems, Life technologies, the Netherlands) using Taqman assays (Applied Biosystems) for 18S RNA (Hs99999901_s1), ADIPOQ (Hs00605917_m1), AR (Hs00171172_m1), BRCA2 (Hs01037421_m1), CRYAB (Hs00157107_m1), CXCL3 (Hs00171061_m1), CXCL12 (Hs00171022_m1), CD55 (Hs00892618_m1), CXCR4 (Hs00237052_m1), DGAT2 (Hs01045913_m1), FGF1 (Hs00265254_m1), FLT3 (Hs00174690_m1), FOXC1 (Hs00559473_s1), IL7 (Hs00174202_m1), IL23A (Hs00900828_g1), IRF8 (Hs00175238_m1), KLB (Hs00545621_m1), LPL (Hs00173425_m1), MAL2 (Hs00294541_m1), NAMPT (Hs00237184_m1), PDGFRA (Hs00998018_m1), PDPN (Hs00366766_m1), PPARγ (Hs01115513_m1), RARRES2 (Hs00414615_g1), RORc (Hs01076122_m1), TGFb1 (Hs00998133_m1), TGFbR1 (Hs00610320_m1) and TNF (Hs99999043_m1). Expression levels of target genes were expressed relative to 18S RNA. An arbitrary cDNA sample was used on each qPCR plate for normalization between different experimental runs.

### Immunohistochemistry

Synovial tissue sections were stained using antibodies against CXCL12 (Monoclonal Mouse IgG1, Clone #79018,Catalog Number: MAB350, R&D Systems, Abingdon, UK), CXCR4 (Monoclonal Mouse IgG2B, Clone #44716; Catalog Number: MAB172, R&D Systems), BRCA2 (Mouse monoclonal IgG1, Clone #5F6, Catalog Number: GTX70123, GeneTex, Irvine, CA) and podoplanin (Monoclonal Rat IgG2a, Clone #NZ-1, Catalog Number: 11-009; Angio Bio, San Diego, CA). The following isotype antibodies were used; isotype mouse IgG1 (Clone #DAK-GO1, Catalog Number: X0931, Dako Cytomation, Glostrup, Denmark) for CXCL12 and BRCA2 stainings, isotype mouse IgG2b (Clone #20116, Catalog Number: MAB004, R&D systems) for CXCR4 staining and isotype rat IgG2a (Clone # eBR2a, Catalog Number: 14-4321, eBioscience) for podoplanin.

Sections (5 μm each) were cut and mounted on StarFrost adhesive glass slides (Knittelgläser, Braunschweig, Germany). Sealed slides were stored at −80°C until further use. Immunohistochemistry was performed using either a two-step (BRCA2 and podoplanin) or three-step (CXCL12 and CXCR4) immunoperoxidase method. Sealed slides containing frozen sections were thawed at room temperature (RT) for 30 minutes, unpacked and air dried for another 20 minutes. Subsequently, sections were fixed in acetone and endogenous peroxidase activity was blocked with 0.3% H_2_O_2_ in 0.1% sodium azide in phosphate buffered saline (PBS) for 20 minutes. After washing in PBS, primary antibodies were incubated overnight at 4 degrees Celsius. As negative control, irrelevant isotype-matched immunoglobulins were applied to the sections instead of the primary antibody. Next day sections were stained using a goat anti-mouse-horseradish peroxidase (HRP)-conjugated antibody (Dako) for BRCA2 and goat anti-rat-HRP-conjugated antibody (Southern Biotech, Birmingham, AL) for podoplanin. For the three-step protocol sections were stained using a goat anti-mouse IgG1-HRP-conjugated antibody (Southern Biotech) for CXCL12 and goat anti-mouse IgG2b-HRP-conjugated antibody (Southern Biotech) after which the intensity was enhanced using a tertiary rabbit anti-goat HRP-conjugated antibody (Southern Biotech). AEC (3-Amino-9-EthylCarbazole; Vector Laboratories, Burlingame, CA) was used as chromogen. Slides were counterstained with Gill’s hematoxylin (Klinipath, Duiven, the Netherlands) and mounted in Kaiser’s glycerol gelatin (Merck, Darmstadt, Germany). Staining was evaluated by microscopy (Leica, Cambridge, UK). Synovial tissue sections collected from patients with early unclassified arthritis (UA), early RA and spondyloarthritis (SpA) patients were used as positive controls for stainings (25). Intensity of staining was analyzed by semi-quantitative image analysis in a blinded fashion by two independent observers.

For the detection of lipid droplets synovial tissue sections were stained using BODIPY 493/503 reagent (4,4-Difluoro-1,3,5,7,8-Pentamethyl-4-Bora-3a,4a-Diaza-s-Indacene, Molecular Probes). Sealed slides containing frozen sections were thawed at RT for 30 minutes, unpacked and air dried for another 20 minutes. Subsequently, sections were fixed in 4% paraformaldehyde and slides were stained one hour at RT with BODIPY (10mg/ml). Slides were counterstained and mounted with VECTASHIELD HardSet Antifade Mounting Medium with DAPI (4’,6-diamidino-2-phenylindole, Vector Laboratories). Staining was evaluated by confocal microscopy (TCS SP8 X, Leica). Per donor, five images were taken using the 20 times oil immersion lens, at 1024×1024 dpi, keeping all settings constant between donors, and images were stored (LIF). Stored images were used for the quantification of BODIPY staining with the full Leica LAS AF software (v2.6.3). The BODIPY signal was converted to a binary signal to allow the calculation of percentage of staining. This was done by setting an arbitrary threshold at 20 gray, resulting in all pixels with an intensity below 20 gray being black (0 gray) and all pixels of 20 gray or higher being fluorescent (255 gray). Only the area of the image representing tissue (based on DAPI staining) was selected for analysis. Of each image the area occupied by tissue (μM^2^), mean value (gray) and maximum value (gray) of BODIPY staining in the tissue area were noted. To calculate the total number of positive pixels and therefore the percentage of staining, the mean value was divided by the maximum value and the resulting value multiplied by 100. Finally by using the area of tissue, the percentage of staining per 100’000 μM^2^ was calculated. The mean value per 100’000 μM^2^ of the five images per donor was used for statistical analysis.

### Statistical analysis

GraphPad Prism Software (V.8.3.0, GraphPad Software, La Jolla, CA) and Statistical Package for the Social Sciences (SPSS) version 26.0 (SPSS, Chicago, IL) were used for statistical analysis. Not normally distributed data are presented as median with IQR. Differences between study groups were analyzed using a Mann Whitney U-test. Categorical data are presented as the number (percent) of subjects and differences between groups were analyzed using the Fisher’s exact test. Cox proportional hazards regression analysis was performed to investigate associations between synovial gene expression profiles and onset of arthritis. Follow-up duration was defined as the time between inclusion in the cohort and the onset of clinically manifest arthritis, or between inclusion and censoring date (for individuals who did not develop arthritis). Due to the low number of events in the Cox proportional hazards analyses, we considered P<0.1 as relevant. Otherwise P values below 0.05 were considered statistically significant.

## Results

Synovial biopsies were collected from in total 67 at risk individuals of whom 17 (25%) developed arthritis after a median follow up time of 15 months. Eventually, 16 at risk individuals developed RA and 1 at risk individual developed osteoarthritis (OA). Demographics of included at risk individuals are listed in Table 1.

**Table 1:** Baseline characteristics of at risk individuals.

|  | Discovery cohort<br>N=13<br>(Agilent arrays) | Total cohort<br>N=67<br>(IHC and/or qPCR*) |
| --- | --- | --- |
| Sex, female (n (%)) | 9 (69) | 48 (72) |
| Age (years) (median (IQR)) | 48 (43 - 53) | 48 (37 – 53) |
| IgM-RF positive (n (%)) | 11 (85) | 37 (55) |
| ACPA positive (n (%)) | 8 (62) | 40 (60) |
| ACPA level ** (kAU/L) (median (IQR)) | 847 (298-2202) | 431 (139 – 1230) |
| IgM-RF and ACPA both pos. (n (%)) | 6 (75) | 13 (19) |
| ESR (mm/hr) (median (IQR)) | 18.0 (8.0-30.5) | 8.5 (2.8 – 18.0)<sup>1</sup> |
| CRP (mg/L) (median (IQR)) | 4.2 (1.8-10.9) | 2.2 (1.0 – 5.9) |
| 68 TJC (n) (median (IQR)) | 5 (2-7) | 2 (0 – 8) |
| 66 SJC (n) (median (IQR)) | 0 | 0 |
| **Developed arthritis (n (%))** | **6 (46)** | **17 (25)<sup>2</sup>** |
| **Follow up time (months) (median (IQR))** | **20 (2 – 44)** | **15 (7 – 35)** |
| No arthritis group: |  |  |
| Follow up time (months) (median (IQR)) | 85 (69 - 86) | 55 (30 – 75) |

*Categorical variables: n (%). Continuous variables (data not normally distributed): median (IQR). ACPA, anticitrullinated protein antibodies; ESR, erythrocyte sedimentation rate; CRP, C-reactive protein; IgM-RF, IgM rheumatoid factor; 68 TJC, tender joint count of 68 joints; 66 SJC, swollen joint count of 66 joints. \*IHC n=67, qPCR n=61, due to quality of RNA. <sup>1</sup>one missing value. <sup>2</sup>16 individuals developed RA and 1 individual developed OA.*

### Microarray profiling reveals gene signatures associated with arthritis development

An explorative genome-wide transcriptional profiling study was performed using synovial tissue obtained from 13 at risk individuals of whom 6 developed RA after a median follow up time of 20 months (IQR 2 – 44). The 7 individuals who did not develop RA had a median follow up time of 85 months (IQR 69 – 86). We used survival analysis to compute the Cox score test for arthritis development for each transcript. Using a False Discovery Rate (FDR) of less than 5% we found that increased expression of 3,151 transcripts correlated with a higher risk of arthritis development, and increased expression of 2,437 transcripts correlated with a lower risk of arthritis development (Supplementary Table 1). Next, two-way hierarchical cluster analyses was used to visualize these significantly differential expressed transcripts between at risk individuals (Figure 1). Except for 1 individual, all pre-RA individuals clustered into the right arm of the dendogram. To interpret the biological relevance of this large list of significantly differentially expressed transcripts we used GSEA analysis to classify these transcripts into canonical pathways (Figure 1). This analysis revealed that higher expressed genes in synovial tissue of pre-RA individuals are involved in several immune response related pathways e.g. T cell and B cell receptor pathways, cytokine and chemokine signaling and antigen processing and presentation. In contrast, a lower expression was observed for genes involved in e.g. extracellular matrix receptor interaction, Wnt-mediated signal transduction and lipid metabolism.

**Figure 1.**
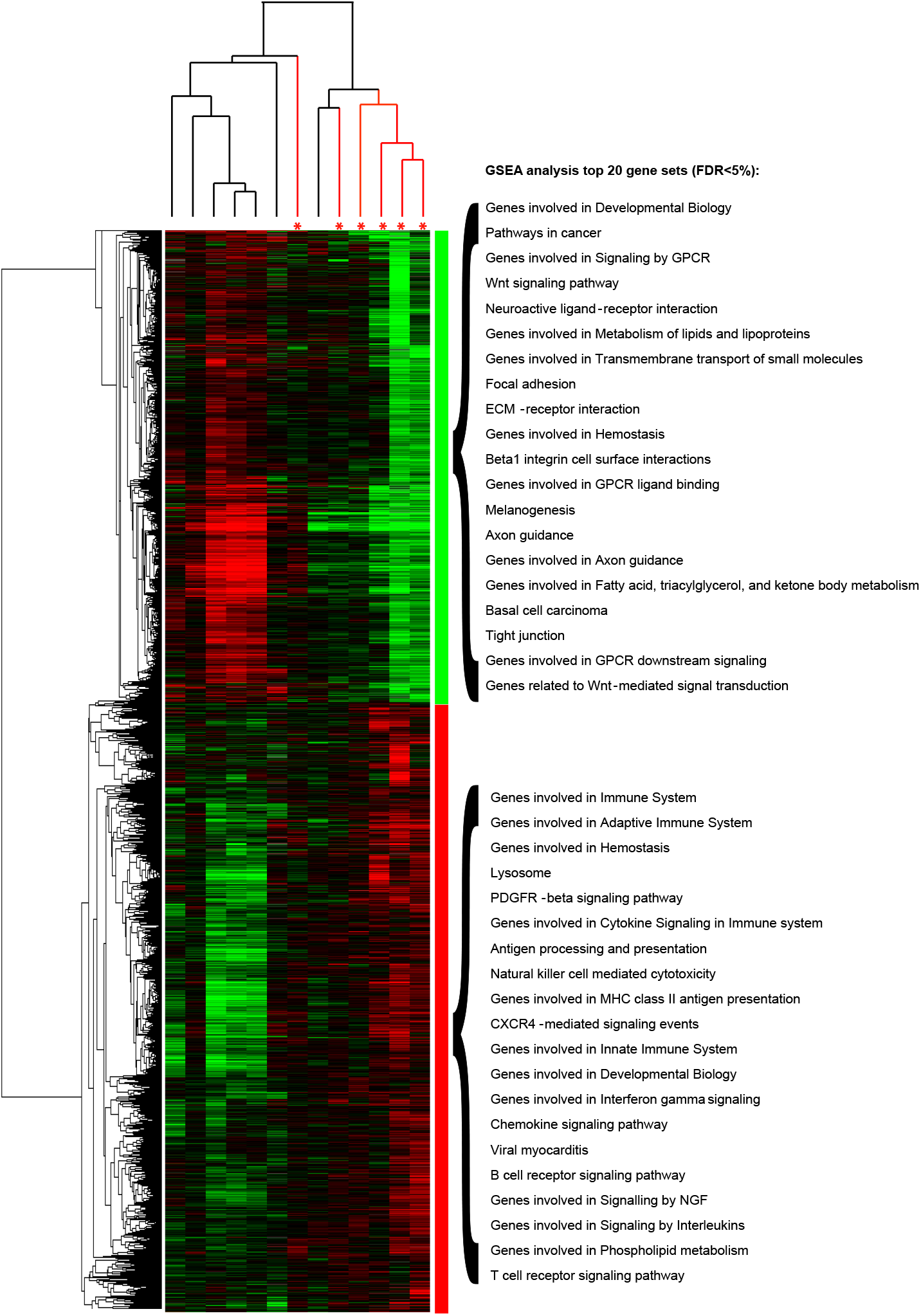
Transcriptional profiles associated with arthritis development. Two-way hierarchical cluster diagram of 5,588 transcripts whose expression levels in synovial tissue were significantly associated with the risk of arthritis development in autoantibody positive arthralgia patients. The cluster diagram is visualized by Treeview. * pre-RA individuals in red lines. Each column represents the data of one array/sample and each row shows the relative expression level of a single transcript for all samples. Red color indicates a relatively higher expression, green color stands for a relatively lower expression and black color indicates that the expression level is equal to the median expression level across all samples. GSEA was used to classify the transcripts into canonical pathways as indicated at the right side of the diagram (showing the top 20 pathways with a FDR<5%).

### Analyses of genes associated with arthritis development in a larger cohort of at risk individuals

Next we aimed to validate our findings in the total cohort of at risk individuals. From 61 at risk individuals (n=16 pre-RA, 1 donor developed OA) high quality RNA could be isolated from synovial biopsies for qPCR analyses. The pre-RA individuals developed arthritis after a median follow time of 15 months (IQR 7 – 35). The remaining 45 at risk individuals had not developed arthritis after a median follow time of 55 months (IQR 30 – 75). From the gene expression profiling study we selected 27 genes whose synovial expression level was significantly associated with arthritis development and whose function has been described to play a role in inflammation or RA. Expression levels of the 27 selected genes were measured by qPCR and their expression levels were used in a two-way hierarchical cluster analysis to visualize association with arthritis development (Figure 2). This cluster diagram classified the at risk individuals into two groups, where pre-RA individuals showed a tendency to cluster together in the left arm of the dendogram (Fisher’s exact test p<0.05). Cox regression analysis was used to test per gene for association with arthritis development. The mRNA expression levels of BRCA2, IRF8, CXCL12, PDGFRA and FLT3 were significantly associated with arthritis development (see Supplementary Table 2). Taken together, differential expression of genes associated with arthritis development could be confirmed in synovial tissues in the total cohort using qPCR analyses.

**Figure 2.**
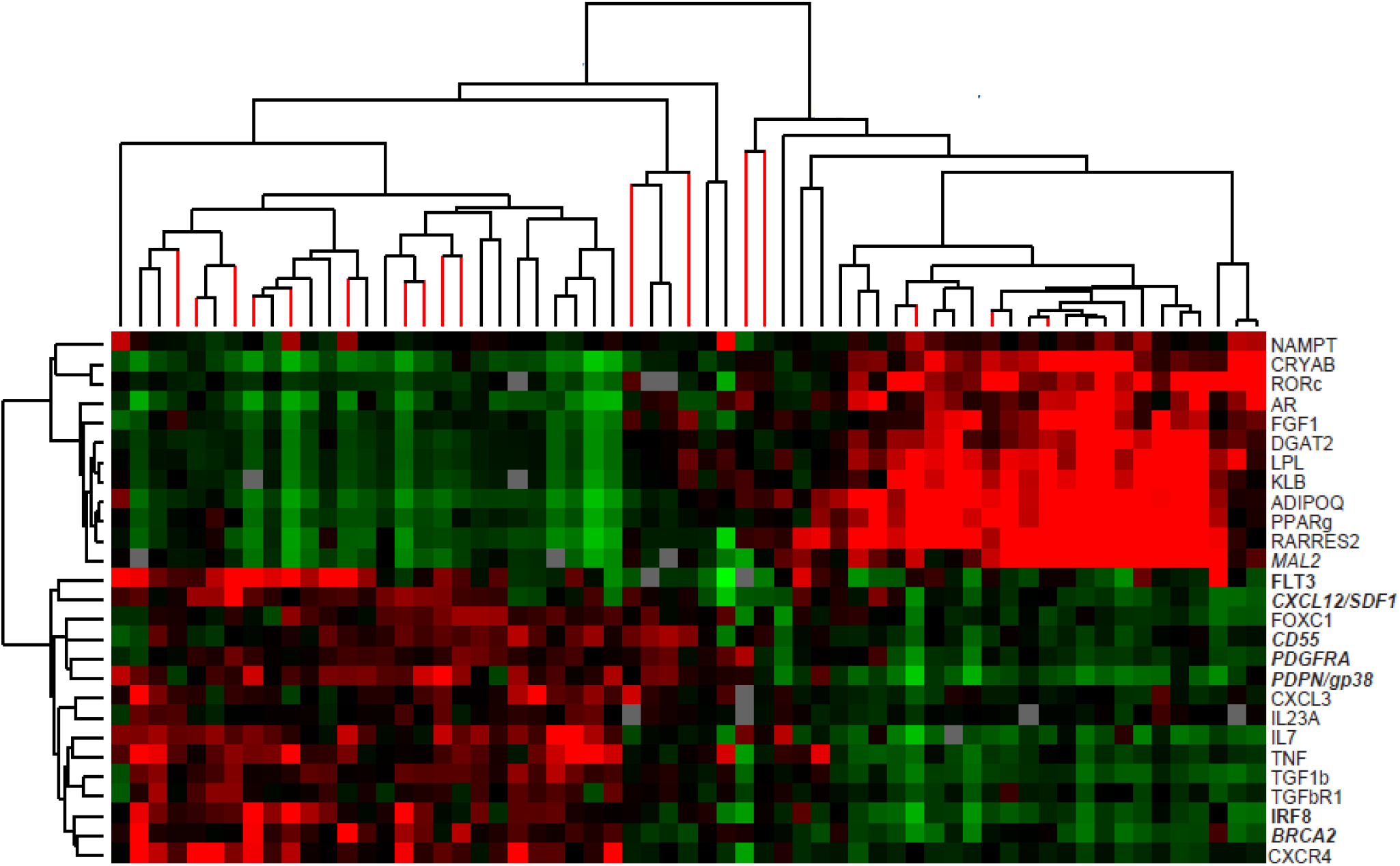
Genes associated with arthritis development confirmed in the total cohort of at risk individuals. Cluster diagram visualizing the expression levels of 27 genes measured using real-time qPCR. Genes in *Italic* are significantly differential expressed between pre-RA individuals and those who did not develop RA (Mann-Whitney U test P<0.05). Genes in **bold** are associated with arthritis development (Cox regression analyses P<0.1). A significant higher proportion of pre-RA individuals is present in the left cluster (Fisher’s exact test P<0.05).

### Detection of CXCL12, CXCR4, BRCA2 and podoplanin in synovial tissues before onset of disease

The results of the GSEA indicates that genes involved in immune responses and cytokine/chemokine signaling are higher expressed in pre-RA synovial tissue, including genes involved in CXCR4-mediated signaling. Using immunohistochemistry we next evaluated in at risk synovial tissues the protein levels for CXCR4 and its ligand CXCL12, BRCA2, a protein involved in cell proliferation (26–28) and podoplanin, described to be expressed on synovial stromal cells (29, 30). The staining intensities for CXCR4 and CXCL12 correlated with each other and although their staining patterns were highly variable between at risk individuals (Figure 3A and 3B), semi-quantitative analyses revealed no significant association with arthritis development (data not shown). BRCA2 staining patterns in synovial tissue from at risk individuals were similar to stainings in synovial tissue from RA patients (Figure 3C) and synovial tissue from pre-RA individuals was more likely to show a positive podoplanin staining in the lining layer compared to synovial tissue from individuals who did not develop RA (Figure 3D). However, semi-quantitative analyses did not show a significant association with arthritis development (data not shown).

**Figure 3.**
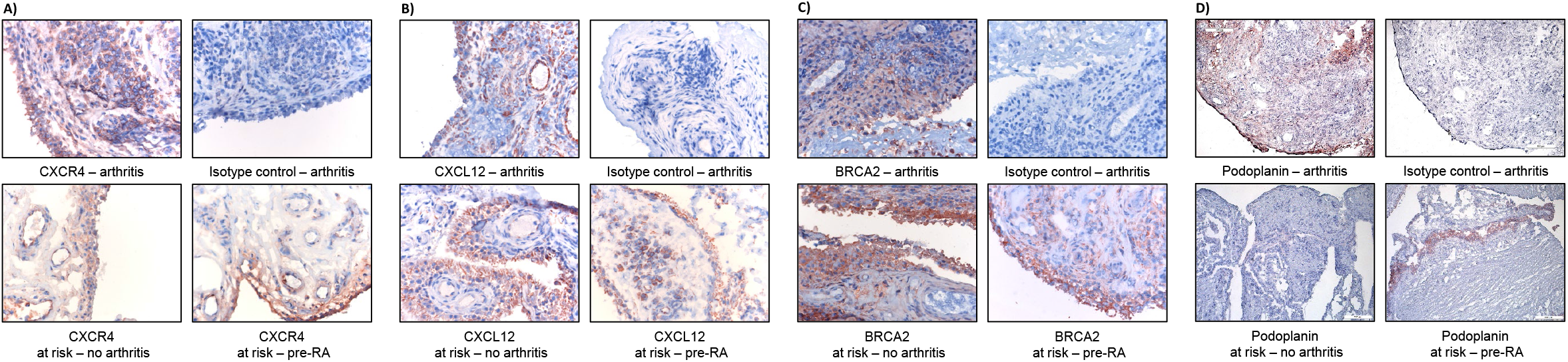
Immunohistochemical analyses of at risk synovium. Detection of CXCR4, 40x magnification (A); CXCL12, 40x magnification (B); BRCA2, 40x magnification (C) and podoplanin, 20x magnification (D) in synovium of pre-RA individuals compared with individuals who did not develop RA. As control, synovium from arthritis patients was stained for CXCR4, CXCL12, BRCA2, podoplanin and isotype controls.

### Differences in lipid droplet formation in synovial tissues

Finally we investigated one of the pathways found to be significantly downregulated in synovial tissue from pre-RA individuals, namely lipid metabolism, as we previously showed an association between increased BMI and arthritis development in this cohort (17). The qPCR analyses already confirmed the downregulated expression of several lipid associated genes: DGAT2, PPARg, RARRES2, ADIPOQ and LPL, in synovial tissues of pre-RA individuals (Figure 2). To visualize and quantify the presence of lipids in the at risk synovium (n=51 of which n=15 pre-RA) we used BODIPY staining which detects lipid droplets. In non-adipocytes, lipid droplets have a protective role as they store fatty acids and convert them into neutral lipids to protect cells from lipotoxicity (31, 32). The observed BODIPY staining pattern was highly variable between synovial tissues as well as within different areas of the synovial tissue sections. Small as well as large lipid droplets could be detected within the synovium (Figure 4A). When we quantified the percentage of BODIPY staining and corrected for synovial tissue volume, we observed a non-significant decrease in lipid droplet formation (P = 0.134) in the pre-RA individuals confirming the microarray data (Figure 4B). Accordingly, at risk individuals with a high (IQR >75) BODIPY staining had a better arthritis-free survival than those with a low (IQR <75) BODIPY staining (Figure 4C; hazard ratio 4.9, 95% confidence interval [95% CI] 0.64-37.10; P = 0.126).

**Figure 4.**
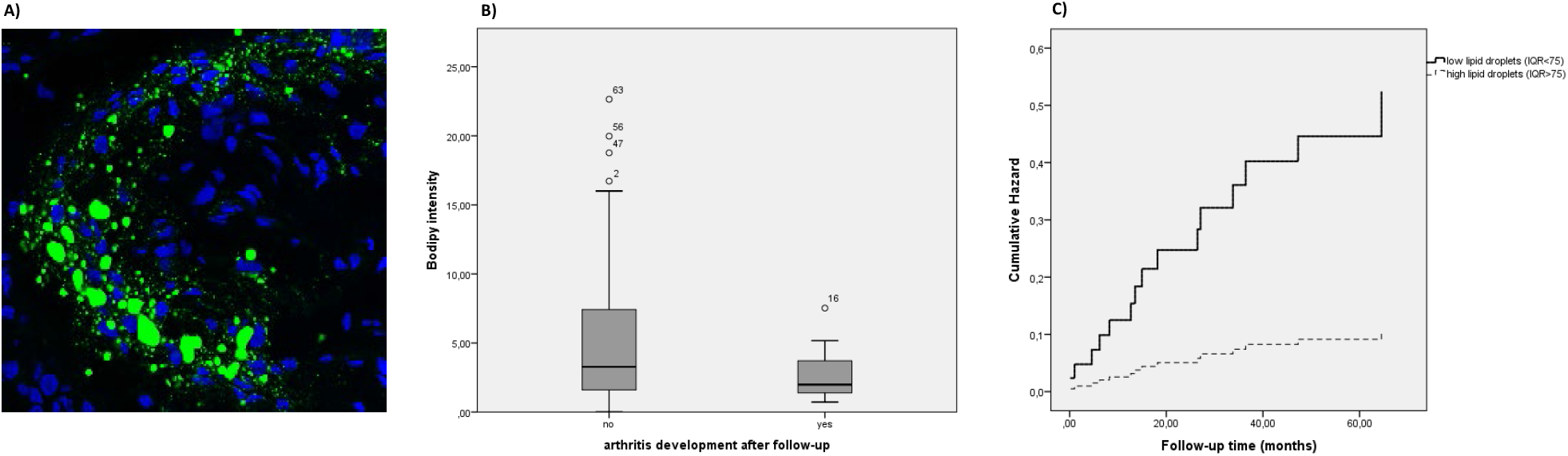
Lipid droplet formation in synovial tissues. Representative BODIPY staining of lipid droplets in the synovium of an at risk individual (A). Quantification of BODIPY staining intensity of synovial tissues of pre-RA individuals and individuals who did not develop arthritis after follow-up (B). Survival curve of at risk individuals with high (IQR>75) synovial BODIPY staining and at risk individuals with low (IQR<75) synovial BODIPY staining (C).

## Discussion

This unique study was set up to investigate if molecular changes are present in the otherwise normal and non-inflammatory synovium of a prospectively followed cohort of individuals at risk of developing RA. Synovial tissues were analyzed using gene expression profiling and microscopy and findings were compared between pre-RA individuals and at risk individuals who did not develop RA after a follow-up of at least 2 years. Results of the explorative expression profiling study revealed gene signatures associated with risk of arthritis development indicating apparent molecular synovial changes before onset of RA.

Previous cross-sectional and prospective studies of autoantibody-positive individuals revealed that systemic autoimmunity, characterized by the presence of disease-specific autoantibodies, precedes the development of clinical arthritis and even synovitis (9, 10). Immunohistochemical analyses of synovial tissue from the same cohort of individuals at risk of developing RA showed that mainly the expression of CD3 and CD8 was associated with arthritis development (10). Our microarray data revealed that synovial tissue of pre-RA individuals display higher expression of genes involved in several immune response-related pathways (e.g. T cell and B cell receptor pathways, cytokine and chemokine signaling and antigen processing and presentation) compared with biopsies of individuals who did not develop RA. In contrast, lower expression was observed for genes involved in e.g. extracellular matrix receptor interaction, Wnt-mediated signal transduction and lipid metabolism. As no overt cellular infiltration is observed during the preclinical phase of RA, these data may reflect changes in resident synovial tissue cells.

The expression levels of 27 differentially expressed genes associated with arthritis development were analyzed in the total cohort of at risk individuals. Two-way hierarchical cluster analysis classified the individuals into two groups, where pre-RA individuals showed a preference to cluster together. GSEA suggests that genes involved in CXCR4-mediated signaling were associated with arthritis development. Immunohistochemistry analyses showed an abundant expression of CXCL12 and CXCR4 in synovial tissues of almost all at risk individuals. CXCR4-CXCL12 binding causes chemokine attraction resulting in the activation and migration of leukocytes during immune and inflammatory responses (33). Our data extends previous studies which show that the number of CXCR4 positive memory T cells in synovial tissue significantly correlate with disease severity (34) and that CXCL12 production by fibroblast-like synoviocytes (FLS) is increased in the synovium of RA patients (35, 36). Our findings suggest that the CXCL12-CXCR4 axis might be activated already during pre-RA. As we previously found no overt cellular infiltration in this preclinical phase of disease (10), the CXCL12 may be derived from stromal cells thereby activating tissue resident T cells. Future functional studies using isolated synovial stromal cells and T cells are needed to address this further.

Interestingly, the mRNA expression level of synovial BRCA2 was significantly associated with arthritis development. Despite abundant presence, we could not confirm this association on protein level. This breast cancer associated gene is mainly detected in proliferating cells (28) and has to our knowledge, not yet been described in synovial tissue. Although the function of BRCA2 is not yet completely defined, BRCA2 is involved in the activation of double-stranded DNA break repair through homologous recombination (37–39) as well as rapidly dividing cells (26–28).

Previous research on synovial tissue and cultured FLS from RA patients showed abundant p53 expression, a tumor suppressor gene activated in response to DNA damage (40). Synovial BRCA2 expression may therefore reflect DNA repair and cellular proliferation in this early phase of RA. Additional research is needed to confirm if genomic instability is already present during pre-RA and if BRCA2 is linked to increased synovial cell proliferation.

Our data suggest an increase in podoplanin expression in synovial tissue before onset of RA. Podoplanin is well-studied in RA and previous studies have shown that it is highly expressed by FLS in synovial tissue of RA patients (29, 41). Research on fibroblast subsets showed that podoplanin expression varies between subsets (30, 42) and that expression on mesenchymal stromal cells can be regulated by inflammatory mediators (42, 43). Furthermore, several studies suggest that podoplanin plays a role in epithelial-to-mesenchymal transition and cellular migration (44–47) and overexpression might be related to a migratory/invasive phenotype of epithelial cells in oral squamous cell carcinomas (44). Overall, our data may suggest that resident mesenchymal stromal cells of pre-RA individuals have an activated phenotype with potentially an increased migratory capacity. This should be explored in future studies.

An unexpected finding was the downregulation of genes involved in lipid metabolism in pre-RA individuals as well as the lower presence of lipid droplets in the synovium. Lipid droplets are important for tissue homeostasis as they store fatty acids to protect non-adipocytes from lipotoxicity (32). Recently, it was shown that lipid metabolism is also important for IL-10 production in T cells (48). The current synovial dataset was used to show that mRNA levels of key enzymes of the cholesterol biosynthesis pathway are significantly associated with arthritis development in at risk individuals suggesting dysregulated IL-10 production (48). In a collagen-induced arthritis mouse model, the inflammatory profile of arthritic mice was linked to defective lipid metabolism (49). In line with our findings, they showed decreased expression of multiple genes related to lipid metabolism including DGAT2, ADIPOQ and LPL (49). Therefore, our data may suggest that lower lipid metabolism is associated with disease progression. In contrast, another study showed accumulation of lipid droplets in synovial T cells of RA patients (50). Further research is needed to investigate the role of synovial cholesterol and lipid metabolism in the pathogenesis of RA.

Unfortunately, the relatively low number of at risk individuals in this study makes it hard to achieve statistical significance. A possible limitation of our study is that biopsies were taken from knee joints, whereas RA usually affects the smaller joints of the hands or feet. However, it is extremely difficult to obtain sufficient synovial tissue from uninflamed synovium from small joints to allow reliable analysis. Nonetheless, studying synovial tissue biopsies from smaller joints may have given similar findings because RA is a systemic and symmetrical disease and it has been demonstrated earlier that synovial inflammation of one joint is usually representative for all inflamed joints (51).

This study describes an exploration of molecular pathways that may help the further understanding of the pathogenesis of RA. Based on these initial data, further research studies to elucidate the importance and role of these pathways in the progression from systemic autoimmunity to the diagnosis of RA can be initiated. As technologies evolve quickly these days, a more comprehensive method to investigate molecular pathways involved in disease progression of RA is single cell RNA-sequencing (scRNA sequencing) although the overall gene coverage per cell is poor. The first studies analyzing RA synovium using scRNA sequencing have been published and revealed that it is possible to distinguish different stages of RA based on single cell gene signatures (52, 53). Using such emerging technologies on synovial tissue samples from at risk individuals would be an interesting next step, though challenging due to the limited number of cells in these uninflamed synovial joints.

In conclusion, our data distinctly shows molecular changes in synovial tissue before clinical manifestation of disease in absence of increased synovial cellular infiltration. Specifically, preclinical synovial changes in immune response genes and lipid metabolism were associated with arthritis development. Validation of the exact role of these pathways in RA may aid the development of preventive RA therapy.

## Authors’s contributions

## Acknowledgements

We thank our study subjects for participating in the study, the AMC arthroscopy team for obtaining synovial tissue biopsies, the AMC KIR lab for sample processing and Iris Simon and Paul Roepman for performing the microarray experiment at Agendia. In addition, we thank the van Leeuwenhoek Centre for Advanced Microscopy facility (AMC) for confocal microscopy. The research leading to these results was funded within the FP7 HEALTH program under the grant agreement FP7-HEALTH-F2-2012-305549, the IMI EU funded project BeTheCure n° 115142, SmartMix subsidy of the Ministry of Economic Affairs of the Netherlands SSM06010, the Dutch Arthritis Foundation n° 11-1-308 and the Dutch Organization for Health Research and Development (ZonMw) Veni n° 916.12.109.

## References

1. Tak PP, Bresnihan B. The pathogenesis and prevention of joint damage in rheumatoid arthritis: advances from synovial biopsy and tissue analysis. Arthritis and rheumatism. 2000;43(12):2619–33.

2. Pollard L, Choy EH, Scott DL. The consequences of rheumatoid arthritis: quality of life measures in the individual patient. Clinical and experimental rheumatology. 2005;23(5 Suppl 39):S43–52.

3. Kyburz D, Gabay C, Michel BA, Finckh A. The long-term impact of early treatment of rheumatoid arthritis on radiographic progression: a population-based cohort study. Rheumatology (Oxford, England). 2011;50(6):1106–10.

4. Ciubotariu E, Gabay C, Finckh A. Joint damage progression in patients with rheumatoid arthritis in clinical remission: do biologics perform better than synthetic antirheumatic drugs? The Journal of rheumatology. 2014;41(8):1576–82.

5. Liu D, Yuan N, Yu G, Song G, Chen Y. Can rheumatoid arthritis ever cease to exist: a review of various therapeutic modalities to maintain drug-free remission? Am J Transl Res. 2017;9(8):3758–75.

6. Chen DY, Lau CS, Elzorkany B, Hsu PN, Praprotnik S, Vasilescu R, et al. Dosing down and then discontinuing biologic therapy in rheumatoid arthritis: a review of the literature. International journal of rheumatic diseases. 2018;21(2):362–72.

7. Gerlag DM, Norris JM, Tak PP. Towards prevention of autoantibody-positive rheumatoid arthritis: from lifestyle modification to preventive treatment. Rheumatology (Oxford, England). 2016;55(4):607–14.

8. Gerlag DM, Safy M, Maijer KI, Tang MW, Tas SW, Starmans-Kool MJF, et al. Effects of B-cell directed therapy on the preclinical stage of rheumatoid arthritis: the PRAIRI study. Annals of the rheumatic diseases. 2019;78(2):179–85.

9. van de Sande MG, de Hair MJ, van der Leij C, Klarenbeek PL, Bos WH, Smith MD, et al. Different stages of rheumatoid arthritis: features of the synovium in the preclinical phase. Annals of the rheumatic diseases. 2011;70(5):772–7.

10. de Hair MJ, van de Sande MG, Ramwadhdoebe TH, Hansson M, Landewe R, van der Leij C, et al. Features of the synovium of individuals at risk of developing rheumatoid arthritis: implications for understanding preclinical rheumatoid arthritis. Arthritis & rheumatology (Hoboken, NJ). 2014;66(3):513–22.

11. Aho K, Heliovaara M, Maatela J, Tuomi T, Palosuo T. Rheumatoid factors antedating clinical rheumatoid arthritis. The Journal of rheumatology. 1991;18(9):1282–4.

12. Rantapaa-Dahlqvist S, de Jong BA, Berglin E, Hallmans G, Wadell G, Stenlund H, et al. Antibodies against cyclic citrullinated peptide and IgA rheumatoid factor predict the development of rheumatoid arthritis. Arthritis and rheumatism. 2003;48(10):2741–9.

13. Nielen MM, van Schaardenburg D, Reesink HW, van de Stadt RJ, van der Horst-Bruinsma IE, de Koning MH, et al. Specific autoantibodies precede the symptoms of rheumatoid arthritis: a study of serial measurements in blood donors. Arthritis and rheumatism. 2004;50(2):380–6.

14. Gerlag DM, Raza K, van Baarsen LG, Brouwer E, Buckley CD, Burmester GR, et al. EULAR recommendations for terminology and research in individuals at risk of rheumatoid arthritis: report from the Study Group for Risk Factors for Rheumatoid Arthritis. Annals of the rheumatic diseases. 2012;71(5):638–41.

15. Tak PP, Doorenspleet ME, de Hair MJH, Klarenbeek PL, van Beers-Tas MH, van Kampen AHC, et al. Dominant B cell receptor clones in peripheral blood predict onset of arthritis in individuals at risk for rheumatoid arthritis. Annals of the rheumatic diseases. 2017;76(11):1924–30.

16. van de Stadt LA, Witte BI, Bos WH, van Schaardenburg D. A prediction rule for the development of arthritis in seropositive arthralgia patients. Annals of the rheumatic diseases. 2013;72(12):1920–6.

17. de Hair MJ, Landewe RB, van de Sande MG, van Schaardenburg D, van Baarsen LG, Gerlag DM, et al. Smoking and overweight determine the likelihood of developing rheumatoid arthritis. Annals of the rheumatic diseases. 2013;72(10):1654–8.

18. Koopman FA, Tang MW, Vermeij J, de Hair MJ, Choi IY, Vervoordeldonk MJ, et al. Autonomic Dysfunction Precedes Development of Rheumatoid Arthritis: A Prospective Cohort Study. EBioMedicine. 2016;6:231–7.

19. Aletaha D, Neogi T, Silman AJ, Funovits J, Felson DT, Bingham CO, 3rd, et al. 2010 Rheumatoid arthritis classification criteria: an American College of Rheumatology/European League Against Rheumatism collaborative initiative. Arthritis and rheumatism. 2010;62(9):2569–81.

20. World Medical Association Declaration of Helsinki: ethical principles for medical research involving human subjects. Jama. 2013;310(20):2191–4.

21. van de Sande MG, Gerlag DM, Lodde BM, van Baarsen LG, Alivernini S, Codullo V, et al. Evaluating antirheumatic treatments using synovial biopsy: a recommendation for standardisation to be used in clinical trials. Annals of the rheumatic diseases. 2011;70(3):423–7.

22. Salazar R, Roepman P, Capella G, Moreno V, Simon I, Dreezen C, et al. Gene expression signature to improve prognosis prediction of stage II and III colorectal cancer. Journal of clinical oncology: official journal of the American Society of Clinical Oncology. 2011;29(1):17–24.

23. Tusher VG, Tibshirani R, Chu G. Significance analysis of microarrays applied to the ionizing radiation response. Proceedings of the National Academy of Sciences of the United States of America. 2001;98(9):5116–21.

24. Eisen MB, Spellman PT, Brown PO, Botstein D. Cluster analysis and display of genome-wide expression patterns. Proceedings of the National Academy of Sciences of the United States of America. 1998;95(25):14863–8.

25. van de Sande MGH, de Launay D, de Hair MJH, García S, van de Sande GPM, Wijbrandts CA, et al. Local Synovial Engagement of Angiogenic TIE-2 Is Associated With the Development of Persistent Erosive Rheumatoid Arthritis in Patients With Early Arthritis. Arthritis & Rheumatism. 2013;65(12):3073–83.

26. Vaughn JP, Cirisano FD, Huper G, Berchuck A, Futreal PA, Marks JR, et al. Cell cycle control of BRCA2. Cancer research. 1996;56(20):4590–4.

27. Rajan JV, Wang M, Marquis ST, Chodosh LA. Brca2 is coordinately regulated with Brca1 during proliferation and differentiation in mammary epithelial cells. Proceedings of the National Academy of Sciences of the United States of America. 1996;93(23):13078–83.

28. Rajan JV, Marquis ST, Gardner HP, Chodosh LA. Developmental expression of Brca2 colocalizes with Brca1 and is associated with proliferation and differentiation in multiple tissues. Developmental biology. 1997;184(2):385–401.

29. Ekwall AK, Eisler T, Anderberg C, Jin C, Karlsson N, Brisslert M, et al. The tumour-associated glycoprotein podoplanin is expressed in fibroblast-like synoviocytes of the hyperplastic synovial lining layer in rheumatoid arthritis. Arthritis research & therapy. 2011;13(2):R40.

30. Mizoguchi F, Slowikowski K, Wei K, Marshall JL, Rao DA, Chang SK, et al. Functionally distinct disease-associated fibroblast subsets in rheumatoid arthritis. Nature communications. 2018;9(1):789-.

31. Martin S, Parton RG. Lipid droplets: a unified view of a dynamic organelle. Nature Reviews Molecular Cell Biology. 2006;7(5):373–8.

32. Bosma M, Kersten S, Hesselink MK, Schrauwen P. Re-evaluating lipotoxic triggers in skeletal muscle: relating intramyocellular lipid metabolism to insulin sensitivity. Progress in lipid research. 2012;51(1):36–49.

33. Buckley CD, Amft N, Bradfield PF, Pilling D, Ross E, Arenzana-Seisdedos F, et al. Persistent Induction of the Chemokine Receptor CXCR4 by TGF-β1 on Synovial T Cells Contributes to Their Accumulation Within the Rheumatoid Synovium. The Journal of Immunology. 2000;165(6):3423–9.

34. Nagafuchi Y, Shoda H, Sumitomo S, Nakachi S, Kato R, Tsuchida Y, et al. Immunophenotyping of rheumatoid arthritis reveals a linkage between HLA-DRB1 genotype, CXCR4 expression on memory CD4+ T cells and disease activity. Scientific Reports. 2016;6(1):29338.

35. Kanbe K, Takagishi K, Chen Q. Stimulation of matrix metalloprotease 3 release from human chondrocytes by the interaction of stromal cell-derived factor 1 and CXC chemokine receptor 4. Arthritis and rheumatism. 2002;46(1):130–7.

36. Kanbe K, Chiba J, Inoue Y, Taguchi M, Yabuki A. SDF-1 and CXCR4 in synovium are associated with disease activity and bone and joint destruction in patients with rheumatoid arthritis treated with golimumab. Modern Rheumatology. 2016;26(1):46–50.

37. Welcsh PL, Owens KN, King MC. Insights into the functions of BRCA1 and BRCA2. Trends in genetics: TIG. 2000;16(2):69–74.

38. Moynahan ME, Pierce AJ, Jasin M. BRCA2 is required for homology-directed repair of chromosomal breaks. Molecular cell. 2001;7(2):263–72.

39. Roy R, Chun J, Powell SN. BRCA1 and BRCA2: different roles in a common pathway of genome protection. Nature reviews Cancer. 2011;12(1):68–78.

40. Firestein GS, Nguyen K, Aupperle KR, Yeo M, Boyle DL, Zvaifler NJ. Apoptosis in rheumatoid arthritis: p53 overexpression in rheumatoid arthritis synovium. Am J Pathol. 1996;149(6):2143–51.

41. Choi IY, Karpus ON, Turner JD, Hardie D, Marshall JL, de Hair MJH, et al. Stromal cell markers are differentially expressed in the synovial tissue of patients with early arthritis. PloS one. 2017;12(8):e0182751.

42. Croft AP, Naylor AJ, Marshall JL, Hardie DL, Zimmermann B, Turner J, et al. Rheumatoid synovial fibroblasts differentiate into distinct subsets in the presence of cytokines and cartilage. Arthritis research & therapy. 2016;18(1):270.

43. Sheriff L, Alanazi A, Ward LSC, Ward C, Munir H, Rayes J, et al. Origin-Specific Adhesive Interactions of Mesenchymal Stem Cells with Platelets Influence Their Behavior After Infusion. Stem cells (Dayton, Ohio). 2018;36(7):1062–74.

44. Martín-Villar E, Scholl FG, Gamallo C, Yurrita MM, Muñoz-Guerra M, Cruces J, et al. Characterization of human PA2.26 antigen (T1α–2, podoplanin), a small membrane mucin induced in oral squamous cell carcinomas. International Journal of Cancer. 2005;113(6):899–910.

45. Astarita JL, Acton SE, Turley SJ. Podoplanin: emerging functions in development, the immune system, and cancer. Front Immunol. 2012;3:283-.

46. Suchanski J, Tejchman A, Zacharski M, Piotrowska A, Grzegrzolka J, Chodaczek G, et al. Podoplanin increases the migration of human fibroblasts and affects the endothelial cell network formation: A possible role for cancer-associated fibroblasts in breast cancer progression. PloS one. 2017;12(9):e0184970.

47. Ward LSC, Sheriff L, Marshall JL, Manning JE, Brill A, Nash GB, et al. Podoplanin regulates the migration of mesenchymal stromal cells and their interaction with platelets. Journal of Cell Science. 2019;132(5):jcs222067.

48. Perucha E, Melchiotti R, Bibby JA, Wu W, Frederiksen KS, Roberts CA, et al. The cholesterol biosynthesis pathway regulates IL-10 expression in human Th1 cells. Nature communications. 2019;10(1):498.

49. Arias de la Rosa I, Escudero-Contreras A, Rodríguez-Cuenca S, Ruiz-Ponce M, Jiménez-Gómez Y, Ruiz-Limón P, et al. Defective glucose and lipid metabolism in rheumatoid arthritis is determined by chronic inflammation in metabolic tissues. Journal of Internal Medicine. 2018;284(1):61–77.

50. Shen Y, Wen Z, Li Y, Matteson EL, Hong J, Goronzy JJ, et al. Metabolic control of the scaffold protein TKS5 in tissue-invasive, proinflammatory T cells. Nature immunology. 2017;18(9):1025–34.

51. Kraan MC, Reece RJ, Smeets TJ, Veale DJ, Emery P, Tak PP. Comparison of synovial tissues from the knee joints and the small joints of rheumatoid arthritis patients: Implications for pathogenesis and evaluation of treatment. Arthritis and rheumatism. 2002;46(8):2034–8.

52. Platzer A, Nussbaumer T, Karonitsch T, Smolen JS, Aletaha D. Analysis of gene expression in rheumatoid arthritis and related conditions offers insights into sex-bias, gene biotypes and co-expression patterns. PloS one. 2019;14(7):e0219698.

53. Lewis MJ, Barnes MR, Blighe K, Goldmann K, Rana S, Hackney JA, et al. Molecular Portraits of Early Rheumatoid Arthritis Identify Clinical and Treatment Response Phenotypes. Cell Rep. 2019;28(9):2455–70.e5.

