## Supplementary table 2 for "Synovial Gene Signatures Associated with the Development of Rheumatoid Arthritis in At Risk Individuals: a Prospective Study"

**Supplementary Table 2 -** mRNA expression levels associated with arthritis development

| Gene | Hazard ratio | P-value | 95% CI | |
|---|---|---|---|---|
| | | | Lower | Upper |
| BRCA2 | 2.375 | 0.019 | 1.149 | 4.906 |
| IRF8 | 3.146 | 0.018 | 1.220 | 8.114 |
| CXCL12/SDF1 | 4.003 | 0.009 | 1.417 | 11.305 |
| FLT3 | 1.936 | 0.029 | 1.072 | 3.496 |
| PDGFRA | 3.582 | 0.042 | 1.046 | 12.264 |
